# Single Cell Foundation Models Evaluation (scFME) for In-Silico Perturbation

**DOI:** 10.1101/2025.09.22.677811

**Authors:** James Boylan, Elizaveta Solovyeva, Theophile Bouiller, Xiong Liu, Sebastian Hoersch, Bulent Ataman, Jeremy Jenkins, Murthy Devarakonda

## Abstract

Foundation models pre-trained on large single-cell RNA atlases offer a compelling alternative to in-vitro experimentation for understanding gene regulatory networks and conducting gene perturbation analyses, with significant implications for target identification. Numerous foundation models have been developed, building upon early efforts such as Geneformer and scGPT. Hyperparameter optimization also results in multiple variants which require comparative analysis. Current benchmarking approaches focus on feature-based assessments or intuitive biological and statistical tasks, which may not align with the models’ training objectives. A recent study proposed a systematic benchmarking framework; however, its scope was limited to pre-trained (zero-shot) models. To address these limitations, we propose Single-Cell Foundation Model Evaluation (scFME)—a systematic method designed to benchmark fine-tuned foundation models for insilico perturbation (ISP). scFME ensures comprehensive and robust assessment by requiring sufficient separation between control and perturbed cells at the outset and by quantifying ISP accuracy against zero and random perturbation baselines. Furthermore, scFME enables exploration of model performance across distinct gene categories, facilitating biological interpretation and functional relevance. Using this framework, we evaluated several commonly used models (and some of their variants) and demonstrated that the methodology effectively characterizes their performance in ISP studies. Our results position scFME as a versatile and rigorous methodology for evaluating and comparing current and future foundation models.

## 1 Introduction

Analyzing gene expression across different cell types and disease states is essential for understanding disease progression and identifying therapeutic targets. Advanced technologies such as single-cell RNA sequencing reveal differences in gene expression at the individual cell level. Leveraging large-scale single-cell RNA sequencing atlases, numerous foundation models based on Transformer architectures—beginning with early and widely cited models such as Geneformer [1] and scGPT [2]—have been proposed to learn latent representations of genes and cells. While these models have demonstrated capabilities across diverse downstream applications, in-silico perturbation (ISP) shows considerable potential and relevance for identifying therapeutic targets.

By using foundation models to predict gene perturbations and their effects on cellular processes, ISP enables rapid, genome-wide target discovery. This approach is efficient, cost-effective, and scalable, offering a viable alternative to traditional experimental methods. However, understanding a model’s in-silico perturbation performance is critical for selecting a suitable model for experiments and building confidence in the results. Robust benchmarking not only facilitates comparison across models, but also helps evaluate hyperparameter configurations and input data preprocessing techniques (e.g., the number of genes selected to represent a cell and relevant cell metadata) for a given model. As a result, benchmarking foundation models has received considerable attention in recent years.

Based on the scope of measurement, benchmarking trends can be categorized as follows: (1) Benchmarking designed to demonstrate specific features for which a model was developed; (2) Benchmarking designed to assess how well a model replicates certain biological functions of cells and their components; (3) Benchmarking designed to assess a model’s performance for a particular task of interest. While these approaches have overlapping attributes, their purposes and assumptions are distinctly different. If one’s interest is in assessing a model for a particular downstream task, such as ISP, then the first two options may be restrictive.

For instance, benchmarking intended to demonstrate design features of a model may not generalize well, as it can lead to different assessments of the same model depending on the features being evaluated, yielding varied outcomes [3]. Such custom benchmarking may include datasets and metrics selected to emphasize the strengths of specific models and applications, which can result in less objective evaluations [4, 5]. Even with standardized datasets and metrics, historical analysis indicates that when benchmarking is conducted by the same group that introduced the new methods, evaluations tend to exaggerate the performance of the latest models through tailored hyperparameters and data processing [3].

Yet another challenge in benchmarking is the expectation of intuitive reproducibility (highlighted by the second category above), which is particularly enticing since we would like the models to be true replicas of biological processes. For example, this approach attempts to benchmark models based on how well they perform on potentially out-of-distribution inputs—for instance, by testing single or double gene perturbations that were not seen during training. A recent study [6] evaluated such predictions and concluded that the performance of often-cited foundation models was not better than simple additive and linear baselines. Another recent study [7] showed that zero-shot ISP prediction performance (i.e., all perturbations are unseen) of Gene-former and scGPT lags behind GEARS, which is trained on perturbation data and tested on a held-out subset of that data. Unfortunately, these approaches do not help us understand model performance for tasks where it can be easily fine-tuned, and this is an important limitation of this category of benchmarking approaches.

While the first category of benchmarking is highly customized for specific model designs (and thus lacks a standardized methodology), the second category focuses exclusively on generalization, which precludes understanding model performance in practical scenarios where they might be effective. To appreciate the subtle relationship between model training and test sets, we consider the manifold hypothesis of deep learning [8]—AI/ML models perform well when test data lies near the low-dimensional manifolds learned during training, as these manifolds capture the intrinsic structure of the data distribution [9]. Recent empirical and theoretical evidence reinforces this observation regarding the performance of deep neural network models [8, 10]. Fine-tuning a pretrained model for a specific perturbed gene can align the learned manifold with the test set and result in good performance. Even minimal training-like interventions, such as chain-of-thought (CoT) prompting, have been shown to guide large language models to activate latent reasoning capabilities [11]. Therefore, benchmarking fine-tuned foundation models on test sets aligned with their training can reveal their strengths and demonstrate their practical usefulness.

In this paper, we propose a systematic methodology that is agnostic to features of any specific foundation model and includes finetuning to evaluate models’ ISP performance under realistic and practical scenarios. The methodology, termed single-cell Foundation Model Evaluation (scFME), builds upon the previous work, especially [7], and uses standard cosine similarity to measure cell state shifts while incorporating rigorous statistical tests for a systematic evaluation procedure. The methodology uses at least one scRNA perturbation dataset for finetuning and evaluation, however, the methodology is independent of the perturbation dataset(s) used. The dataset is expected to contain scRNA sequencing data for control cells and perturbed cells of each perturbed gene. The key characteristics of the scFME framework are:

- The perturbed genes used as ground truth are “qualified” based on their gene expression to ensure viable assessment
- The model under evaluation is finetuned to classify control and perturbed cells to facilitate learning a low-rank manifold for improved ISP
- The framework ensures sufficient separation (an adjustable parameter) between the finetuned control and perturbed cell embeddings in the latent space, which allows meaningful measurement of ISP effects
- Cosine-Shift (the difference between the cosine similarities of control-perturbed and ISP-perturbed cells) is used to measure the model’s ability to reproduce in-vitro perturbations
- The number of genes for which the model statistically outperforms two baselines, the zero cosine-shift and random perturbations cosine-shift, represents its overall ISP performance
- By stratifying a model’s successful ISP performance in terms of the biological functions of the perturbed genes, the framework attempts to provide further insights into a model’s representation of cellular dynamics

The single-cell Perturb-seq perturbation dataset from Weissman Labs [12], specifically the K562 Essential dataset, was used to demonstrate the proposed methodology. Three Geneformer models, two scGPT variants, and GenePT were assessed to validate the methodology’s capability. As mentioned earlier, although the Perturb-seq dataset was used as the primary evaluation dataset here, the framework is broadly applicable to other perturbation datasets. It should be noted that the models are fine tuned for cell classification, which is always possible with the datasets available for target identification, rather than requiring perturbation datasets for finetuning.

Our results show that scFME enables systematic evaluation of foundation models for ISP at the cellular level within the context of the selected evaluation dataset. This study establishes a robust framework that is neither exclusively emphasizes model features nor limits understanding to out-of-distribution scenarios only (i.e., not necessarily aligned on a low-rank manifold).

Of the foundation models we studied with the framework, the second generation Geneformer trained on additional cancer cell atlas outperforms the other models, including the first generation Geneformer, the two scGPT models and GenePT.

Larger models pretrained on bigger cell atlases perform better, supporting the scaling laws. One fundamental difference among the model families is the cell representation in the model input, and so we hypothesize that our framework is showing the effects of the representational differences for the ISP task. Further studies are needed to examine if performance of these model-families changes with increased pretraining data and/or hyperparameter optimization. Studies are also needed to see the impact of larger perturbation datasets. The proposed framework is an effective approach to understand the impact of these considerations.

## 2 Results

### 2.1 scFME enables benchmarking ISP performance

The key result of this work is a robust benchmarking framework for understanding single cell foundation models’ performance for in-silico perturbation. The process for benchmarking, one or more models, starts with selecting a perturbation data set with the broadest coverage of perturbed genes and the largest number of cells perturbed per gene. We used the K562 Perturb-seq dataset from Weissman lab [12] The number of perturbed genes were 2,057, yielding 310,385 perturbed cells, with an average of 117 cells per perturbed gene. There were 10,305 control cells in the dataset. An application of the framework follows these steps:

#### 1. Selecting ground truth genes based on gene expression

First, genes with significant expression changes after perturbation in the original in-vitro experiment are identified in the dataset. Each gene perturbation is assessed based on two criteria: (a) whether its log fold change in perturbed cells relative to control cells exceeds a defined threshold with an adjusted p-value < 0.05, and (b) whether it induces substantial transcriptomic changes, as evidenced by a sufficient number of successfully perturbed cells and a statistically significant E-distance [13] (see Methods) compared to controls. Only the perturbed genes meeting both the criteria are chosen as the ground truth genes for further experiments, ensuring effective knockdown and distinctive transcriptional changes.

#### 2. Fine tuning models for each ground truth gene

For each ground truth gene, perturbed cells are paired with control cells and are used to finetune the model under consideration to predict control vs. perturbed cells. The standard finetuning approach is used, i.e. a stratified random sample from the clusters is used for training, and a held-out test set is used to measure precision, recall, and F1 measures. Thus, the framework creates as many finetuned models as the number of ground truth genes.

#### 3. Test of control and target cell clusters separation in the latent space

This step checks whether control and target cell clusters for a perturbed gene are 5 clearly distinct in the finetuned model’s latent space, using silhouette score as the metric. Clear separation is essential; without it, analysis of ISP performance using cosine shift would not yield interpretable metrics, and so this test ensures valid conditions for further ISP evaluation. Only the perturbed genes that achieve silhouette score greater than 0.3 are used in subsequent steps, however, a different threshold may be used for more distinct or less distinct separation. We determined the threshold through trial and error. For a given model, *GPSS* _0.3_ is used as a performance metric. It represents the number of **g**ene **p**erturbations with a **s**ilhouette **s**core above the threshold of 0.3

#### 4. In-silico perturbation and cosine distances determination

For each gene that met the silhouette score threshold, ISP is conducted using corresponding finetuned model, which creates in-silico representations of perturbed cells (“virtually perturbed cells”). For this study we only considered down regulation, but up regulation can also be used in a similar way. Down regulation is implemented according to the model design and authors recommendation. After ISP is conducted, cosine distances are calculated between in-silico perturbed and in-vivo perturbed (target) cells, and between control and target cells. While the number of control and in-silico perturbed cells are always equal (for a perturbated gene), the number of target cells are different from control/ISP cells, and so the centroid of the target cells is used for calculating cosine distances, see Methods for details. Baseline comparisons, along with relevant statistical tests, are conducted using the differences between these cosine distances to determine if the model made successful ISP predictions as described below.

#### 5. ISP performance analysis using baselines and statistical tests

We use two baselines and relevant statistical tests to evaluate a model’s ISP performance for a perturbed gene. The key metric, cosine shift, is measured as the difference between the cosine similarity of ISP cells vs. target cells and control cells vs. target cells; this value ranges from -2 (the worst case) to 2.0 (the best case). We apply a one-sided Wilcoxon signed-rank test to check if the cosine shift exceeds zero (the first baseline) and is statistically significant. Second, we separately generate in-silico perturbations of 100 random genes and use their cosine shifts as the random baseline threshold, against which a one-sided Wilcoxon ranksum test determines if a gene’s cosine shift surpasses this baseline. The number of genes passing these two baselines serve as the ISP performance measures for a foundation model. We introduce two metrics to measure success; *GPSI* _zero_ which is the number of **g**ene **p**erturbations statistically **s**hifted **i**n-silico relative to the zero baseline, and *GPSI* _random_ which is the number of **g**ene **p**erturbations statistically **s**hifted **i**n-silico relative to the random baseline.

#### 6. Evaluating model accuracy across biological functional annotations

An additional step in our methodology involves exploring each model’s ISP performance across gene categories, which are defined based on biological functions using gene set and pathway analysis databases. This analysis demonstrates how various models identify perturbation within — both overlapping and unique — functional categories such as ribosomal genes and translation initiation factors. Such comparisons can elucidate the relative strengths and limitations of each model in detecting specific biological perturbations.

We applied the scFME framework and identified 224 ground truth genes, see Methods. Foundation models across three families were selected for evaluation: Geneformer (three variants), scGPT (two variants), and GenePT.

These model families use distinctly different cell representations. Geneformer uses rank ordering of genes based on gene expression normalized across cells, while scGPT maps gene expression to expression-value bins and combines encoding of bins and gene ids. GenePT utilizes Large Language Model (LLM) embeddings from textual gene descriptions. scFME distinguishes performance differences resulting from these different input representations, pretraining atlas sizes, and model sizes.

Each model-variant, except GenePT (which was not designed to be finetuned), was individually finetuned on the 224 genes using the K562 dataset. Finetuning for the task of classifying control and perturbed cell types yielded 224 separate fine-tuned versions of each model. The precision, recall, and F1 measures on the held-out test sets were determined across the genes and models. The subsequent steps of the framework, i.e. silhouette score separation in the latent space, ISP and cosine distances determination, cosine shift calculation and baseline comparisons, and further analysis using biological functional categories, were conducted on the above-mentioned models. These experiments demonstrated the viability of the framework, and the results from these experiments are further described below.

### 2.2 ISP performance differences of foundation models

The scFME framework identified performance differences among the single cell foundation model families for ISP (Table 1). We evaluated the first generation Gene-former pretrained on 30M cell atlas, with 2K input tokens and about 10M parameter model, and the second generation Geneformer using 95M and 104M training atlases, with 4K input tokens and 38M parameters. We evaluated the scGPT model that was trained on a 33M cell atlas, with 3000 input tokens (plus the CLS token) and 51.3M model parameters, denoted by scGPT-RS. A second scGPT model was evaluated by selecting 3,000 highly variable genes as the preprocessing step, denoted by scGPT-HVG, see Methods.

**Table 1.**
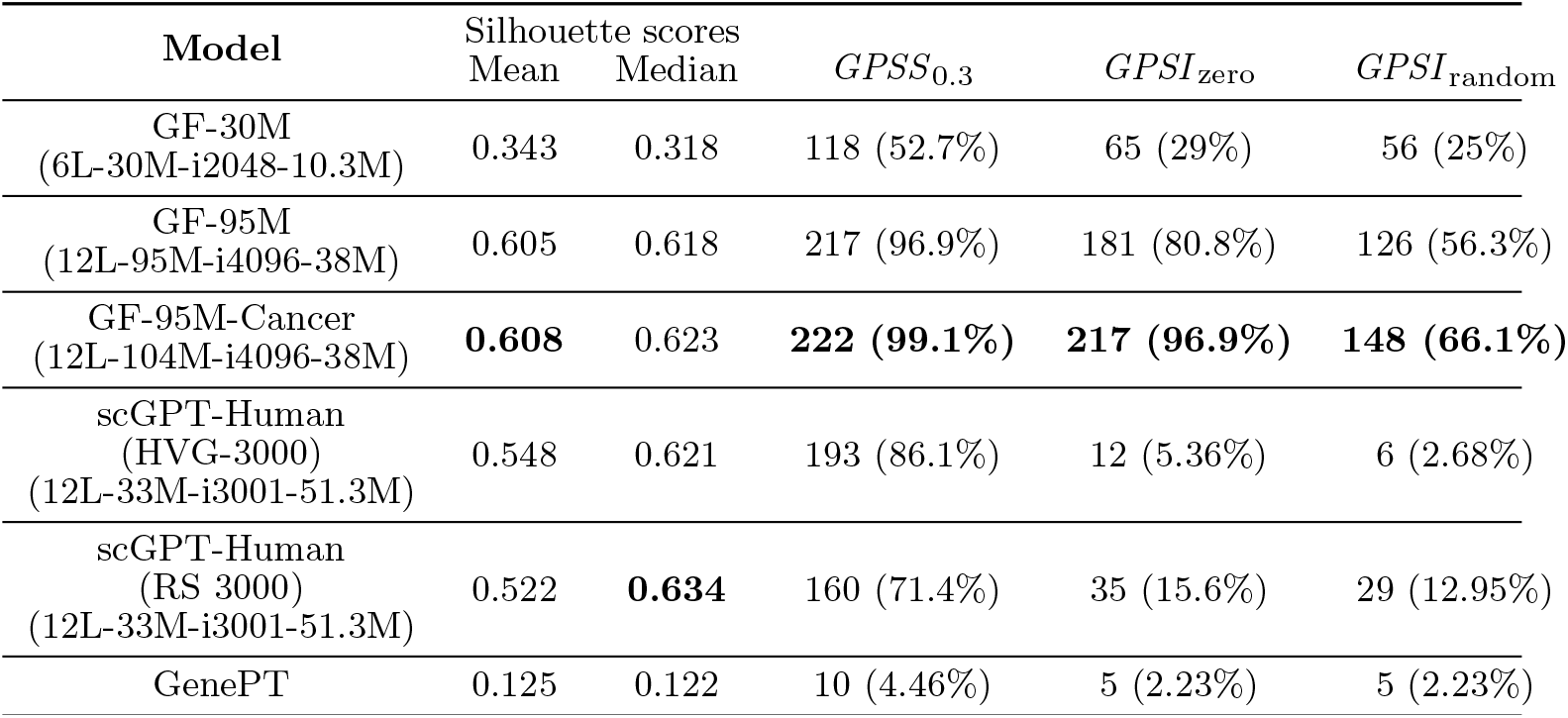
Summary of silhouette scores and ISP success rates for seven models. Columns report mean and median silhouette scores, number of genes passing the silhouette threshold ( ≥0.3), and number of genes with positive cosine shift exceeding zero and random baselines.

**Table 2.**
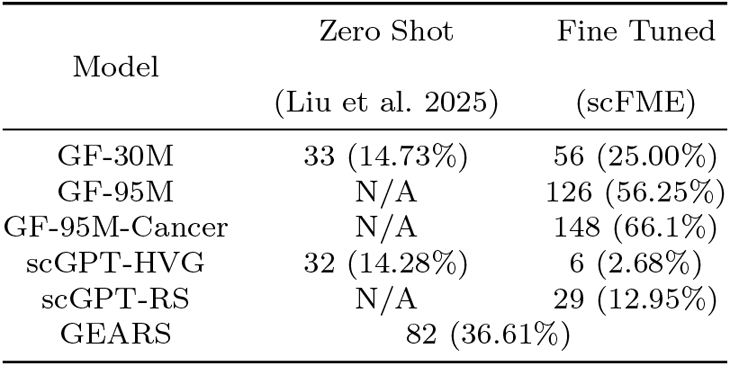
Number and percentage of ground truth genes successfully predicted by each model under zero-shot (from [7]) and fine-tuned (scFME) conditions.

scFME benchmarking showed that Geneformer-95M-Cancer, which was the 95M Geneformer that was additionally pretrained using cancer-specific data (a total of 104M cells), demonstrated consistently high performance across most metrics. It achieved a silhouette score ≥0.3 for 222 of 224 genes (99.1%) with the best mean silhouette score of 0.608 and passed the ISP cosine shift test for 217 genes (96.9%). Notably, 148 genes (66.1%) exhibited a statistically significant shift greater than random, indicating robust alignment between in-silico and in-vitro perturbation states.

In contrast, scGPT-RS achieved the highest median silhouette score (0.634), suggesting strong latent space separation overall. However, only 160 genes (71.4%) passed the silhouette score threshold, and 29 genes (12.95%) passed the ISP test with a shift greater than random. While the scGPT-RS model we evaluated captures cell state structure effectively, its perturbation predictions are less biologically aligned. GenePT underperformed across all metrics, with only 10 genes (4.46%) passing the silhouette score threshold and just 5 genes (2.23%) showing significant ISP shifts. It should be noted that these results are for the model configurations we used, and further studies are needed to understand the performance differences of the models under hyperparameter optimized conditions.

### 2.3 The scaling law and domain-specific pretraining

Performance comparison of the three variants of the Geneformer model family revealed how model size and training data influence in-silico perturbation performance.

Across all the metrics, performance improved with model scale. Compared to Geneformer-30M, which was pretrained on 30 million healthy cells, the 95M model, trained on over three times more cells and has larger model size and increased input length, demonstrated substantial gains in both separation accuracy and ISP performance. The Geneformer-95M-Cancer model, which incorporated an additional 14 million cancer-specific cells into its pretraining corpus, achieved the highest performance overall—surpassing both its variants. It should be noted that the K562 perturbation dataset was generated from a cancer cell line.

The observed performance improvement across the Geneformer variants supports the hypothesis that the larger models trained with biologically aligned pretraining data are better equipped to capture the nuanced transcriptomic shifts induced by gene perturbations. By incorporating cancer-specific data, Geneformer-95M-Cancer likely learned manifolds more representative of the K562 cell line used in evaluation, thereby improving alignment between in-silico and in-vitro perturbation states.

Future iterations of foundation models for single-cell biology may benefit from targeted scaling strategies—such as increasing model size, input dimensionality, and pretraining on disease-relevant datasets—to optimize performance for specific biological tasks. These findings highlight the utility of the scFME framework not only as a benchmarking tool but also as a guide for model development.

### 2.4 Determinants of successful in-silico perturbations

To identify the factors underpinning successful in-silico perturbations, we further analyzed factors that might have influenced in-silico perturbation accuracy, for the Geneformer model family. As mentioned earlier, ISP success is defined as cosine shift greater than the random baseline, with statistical significance of adjusted p-value < 0.05.

We analyzed three types of factors: in-vitro perturbation strength, fine-tuning performance, and ISP-specific metrics, see Figure 2. Among all the factors, the log2 fold change of the perturbed gene emerged as a strong predictor of ISP success across all Geneformer variants. Perturbations with pronounced transcriptional knockdown were likely to yield successful ISP predictions, suggesting that models are more effective when the biological signal is robust, although it is not exclusive.

**Fig. 1.**
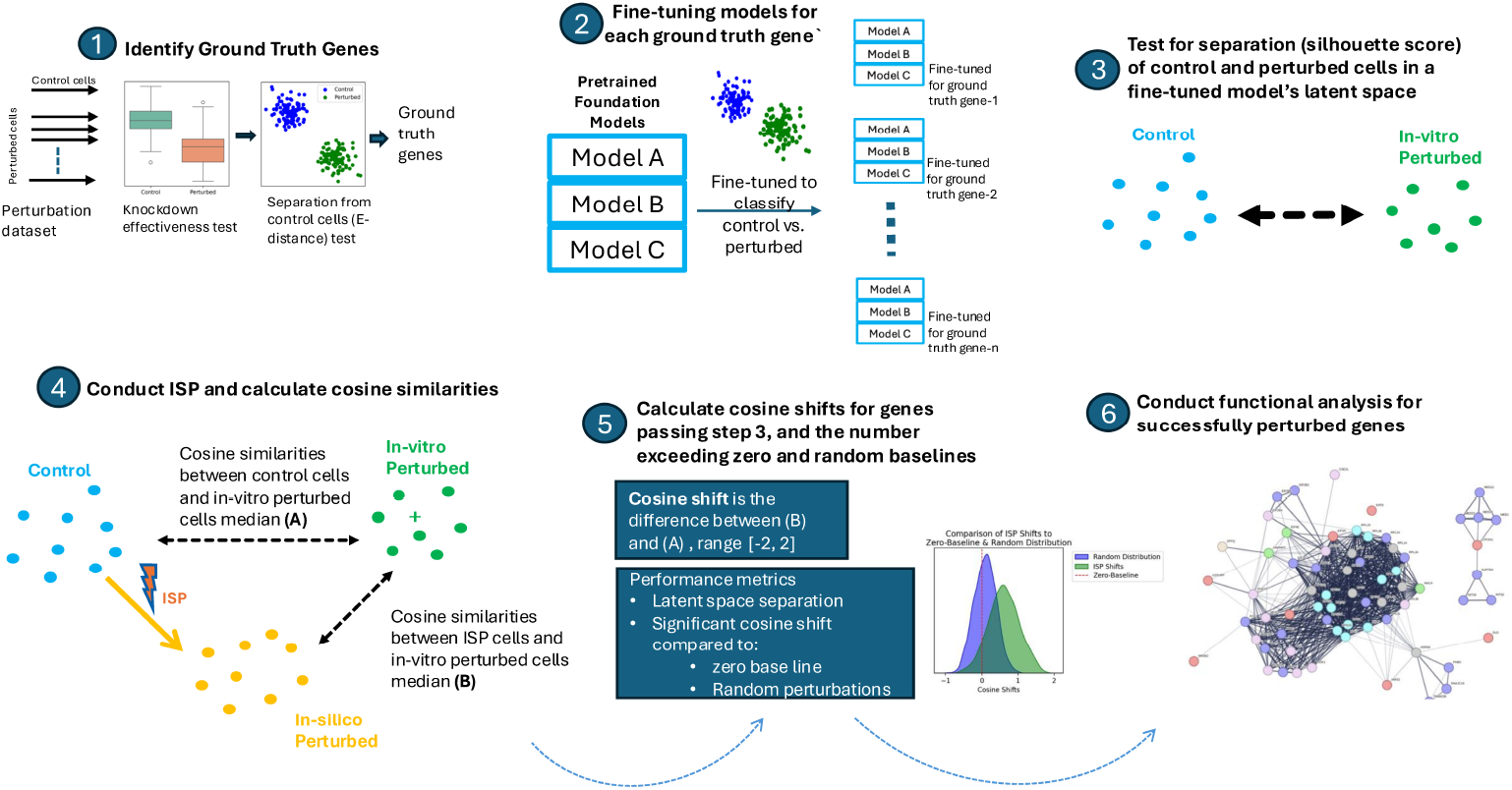
Schematic representation of the six-step scFME pipeline for evaluating foundation models on in-silico perturbation (ISP) tasks. The workflow includes ground truth gene selection, model finetuning, latent space separation assessment, ISP generation, cosine shift analysis against baselines, and functional category evaluation.

**Fig. 2.**
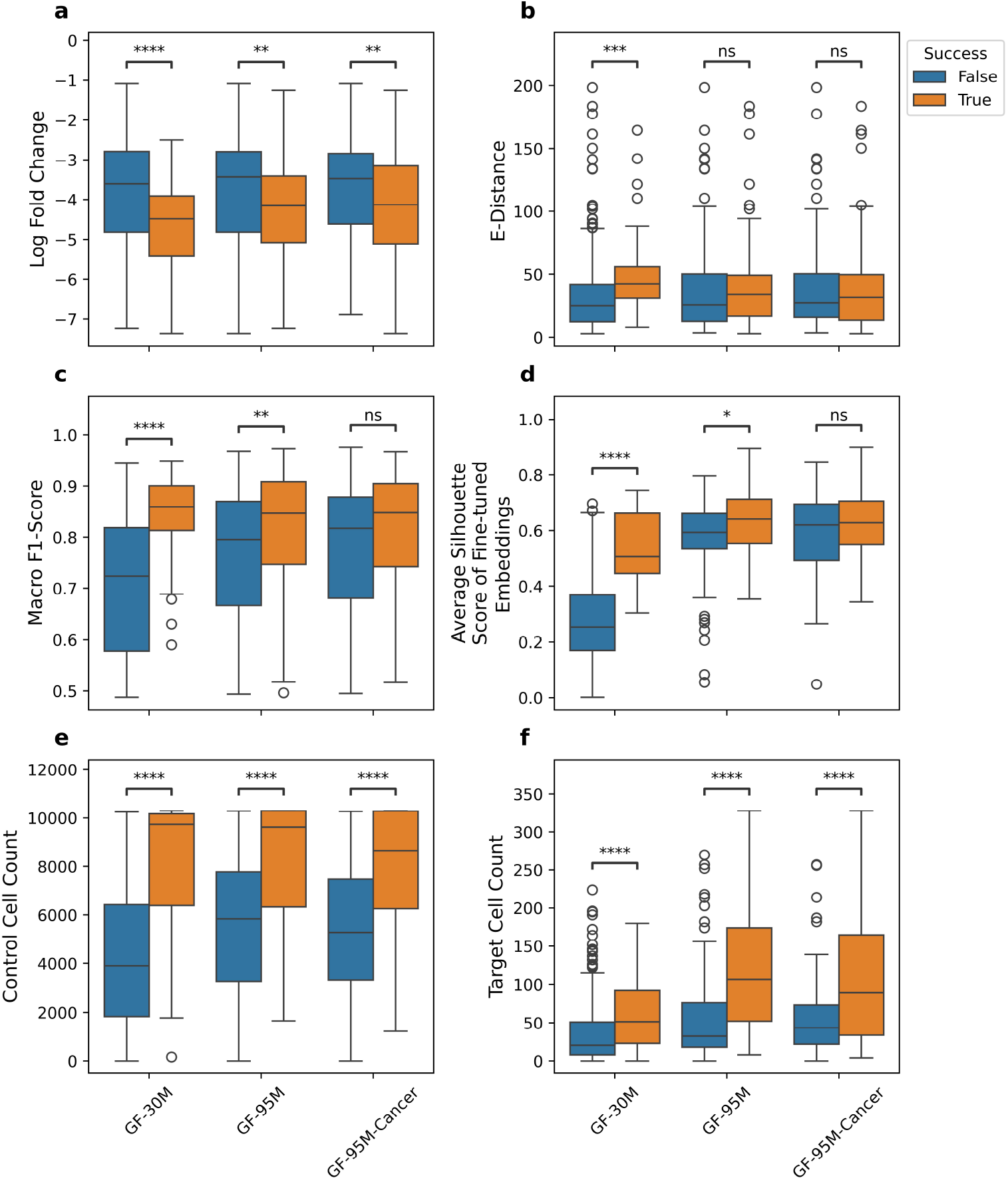
Correlation of ISP success with biological and model-specific factors across Geneformer variants. Panels show associations with (a) log2 fold change of perturbed genes, (b) E-distance between control and perturbed cells gene expression, (c) fine-tuning macro F1-score, (d) silhouette score, (e) and (f) show the number of cells with the ground truth gene in the cell representation for control and perturbed cells respectively.

E-distance, a measure of transcriptomic divergence between the control and perturbed cells, based on the raw gene expression, was significantly associated with ISP success only in the Geneformer-30M model (adjusted p-value ≤1*e* −3), but not in the larger 95M variants. This suggests that the smaller model may be more sensitive to the raw transcriptomic differences, whereas the larger models may rely more heavily on targeted perturbation signals.

Fine-tuning performance, as measured by macro F1-score and silhouette score, also correlated with ISP success. Geneformer-30M exhibited the strongest association, with successful perturbations achieving a median F1-score of 0.86 versus 0.72 for unsuccessful ones (adjusted p-value ≤ 1*e* − 4). Geneformer-95M showed a smaller but still significant difference (0.85 vs. 0.80; adjusted p-value ≤ 1*e* − 2), while Geneformer-95M-Cancer showed no significant difference, potentially reflecting the benefits of domain-specific pretraining in stabilizing performance across perturbation strengths. Latent space separation, quantified by silhouette scores, followed a similar trend. Geneformer-30M again showed the most pronounced difference between successful and unsuccessful perturbations (adjusted p-value ≤ 1*e* − 4), followed by Geneformer-95M (adjusted p-value ≤ 5*e* − 2), with no significant difference observed for the 95M-Cancer model, the cancer-adapted variant.

Finally, we examined the influence of cell representation density. Across all models, successful perturbations were associated with higher counts of both control and target cells (adjusted p-value ≤ 1*e* − 4), indicating that perturbations affecting more prevalent genes are more likely to be captured accurately. Notably, the Geneformer-95M variants had higher median cell counts than Geneformer-30M, likely due to their larger input size, which may enhance their ability to model perturbations in more frequently expressed genes.

Together, these findings suggest that ISP success is influenced by both biological signal strength and model-specific factors, including sensitivity to fine-tuning and latent space structure. The scFME framework thus provides a principled approach for dissecting the conditions under which foundation models most effectively simulate gene perturbations.

### 2.5 ISP performance and differential gene expression (DGE)

To further explore the role of DGE in ISP prediction for various models, we compared the level of ground truth gene down-regulation (fold change) and successful ISP prediction, see Figure 3a. Genes, whose ISP was successfully predicted by one or more models, demonstrated significantly lower fold change values (i.e. larger down regulation) compared to the genes, whose ISP is not successfully predicted by any model. We only considered the Geneformer models and scGPT-RS for this analysis; the other models were excluded due to the low number of successful ground truth genes.

**Fig. 3.**
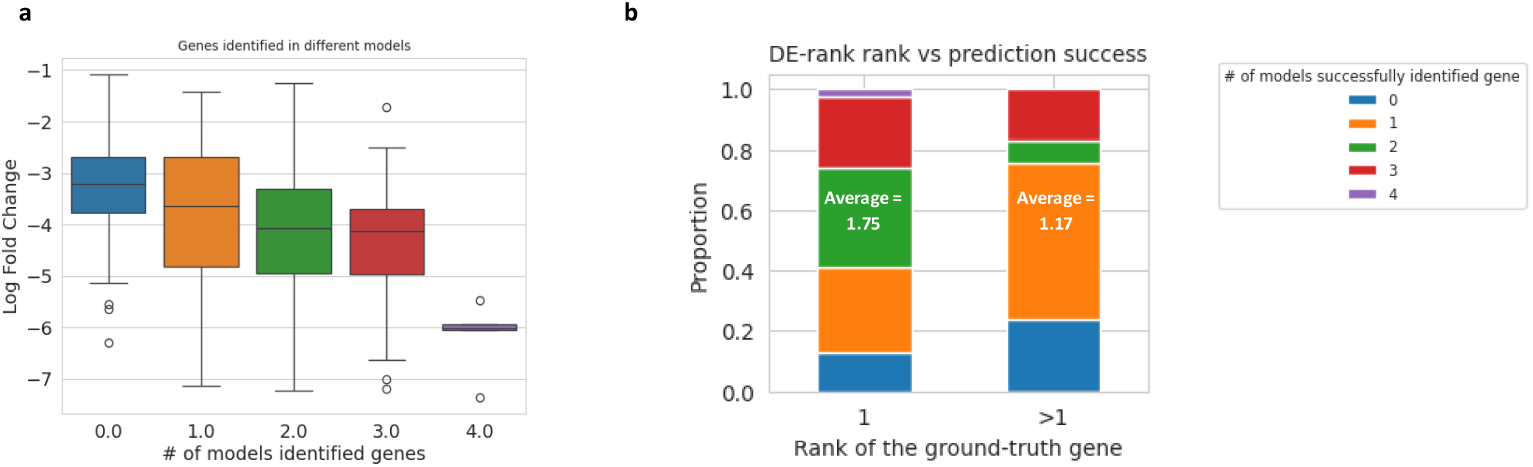
(a) Distribution of log2 fold change for genes successfully predicted by one or more models versus those not predicted. (b) ISP success rate stratified by differential expression (DE) rank, showing higher accuracy for genes ranked as most down-regulated.

**Fig. 4.**
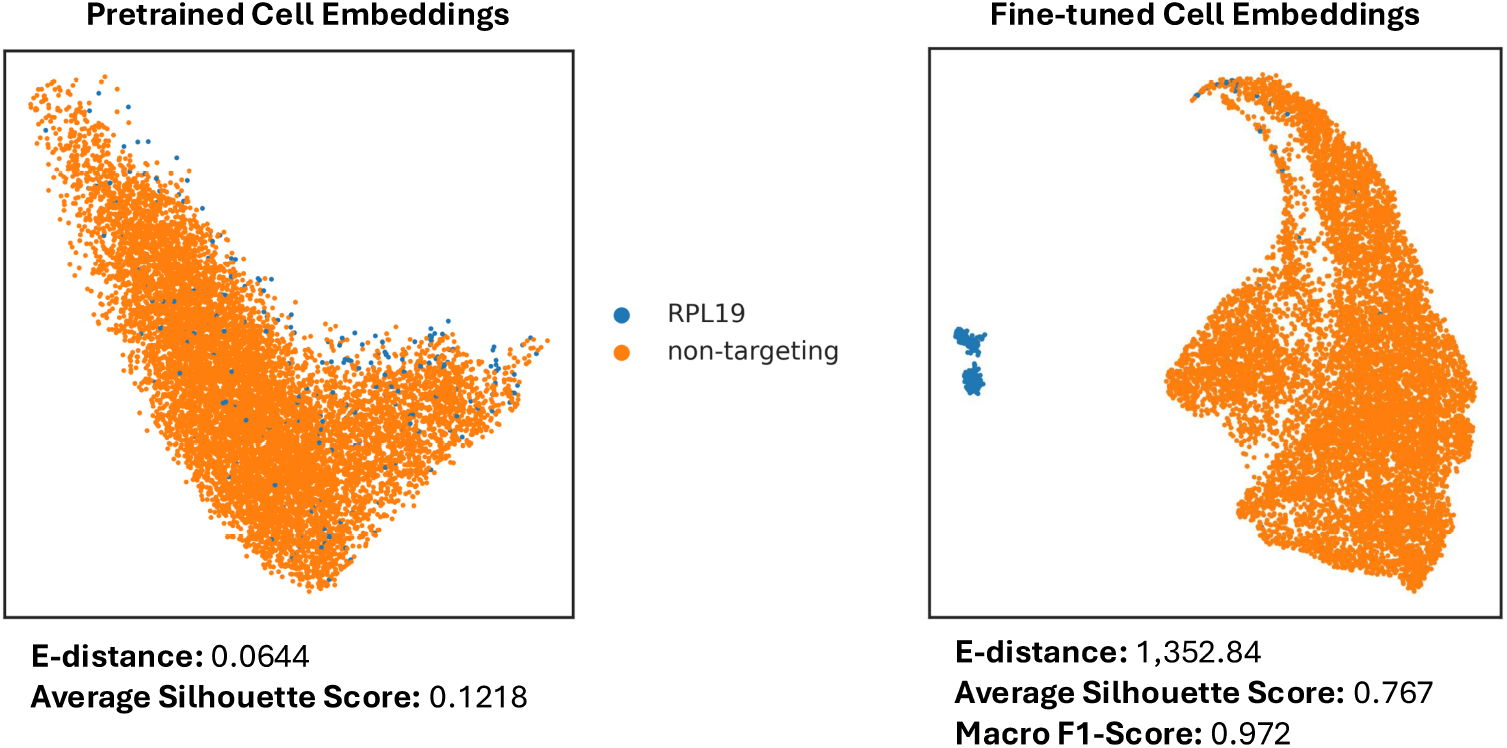
UMAP of cell embeddings for one of the ground truth genes, RPL19, using the Geneformer-95M model. The left plot shows the cell embeddings for Control and RPL19 perturbed cells from the pretrained Geneformer-95M model, indicating very little separation between the two cell states (Silhouette score of 0.1218). The right plot shows the cell embeddings for Control and RPL19 perturbed from the fine-tuned Geneformer-95M model, indicating a high degree of separation between the two cell states (Silhouette score 0.767)

Furthermore, we wanted to see if ISP is more accurate when the perturbed gene is the most downregulated gene. First, we used the t-test from scanpy package to identify the most downregulated genes using the *π*-statistic [14], which resulted in 195 of the 224 ground truth genes being identified as the most downregulated.

We assigned DE (differential expression) rank of 1 to the 195 genes and greater than 1 to the remaining. As seen in Figure 3b, more models successfully predicted ISP when the gene’s DE rank is 1. On the other hand, genes with DE rank higher than one generally showed lower ISP performance across all models, with Geneformer-95M-Cancer showing the best results (15 out of 29 genes).

To further evaluate the relationship between the DE rank of a gene and ISP performance of the GF-95M-Cancer model for the gene, we created a separate list of 122 additional genes. The genes in the list met the following criteria: (1) each originating from a different biological pathway and (2) with a DE rank > 1. The selection criteria were the same as for the original 224 genes, except for the E-distance cut-off. Among these 122 genes, ISP prediction was successful for 28 (23%). This result suggests that the model’s ISP prediction may be less accurate for genes that are not the most differentially expressed, highlighting an area for potential improvement. The scFME framework is useful in identifying such areas of further investigation. This result may also indicate a limitation of the PerturbSeq dataset for evaluating model performance on real-world datasets, where the causal gene is often not the most differentially expressed gene.

### 2.6 Fine-tuning enhances ISP performance

Since we hypothesized finetuning models would enhance ISP performance by aligning the test set with the training manifolds, we wanted to quantify the impact of finetuning on ISP performance. We compared foundation models evaluated using the scFME framework against their zero-shot counterparts previously benchmarked using the scISP methodology [7]. While zero-shot Geneformer-30M and scGPT models were evaluated without exposure to perturbation data, scFME finetuned each model for 224 ground truth genes using a part of the perturbation data, enabling direct comparison of performance gains attributable to finetuning.

Finetuning led to improvements in ISP accuracy for Geneformer-30M, increasing from 14.73% for the zero-shot model to 25%. This improvement reflects enhanced alignment between in-silico and in-vitro perturbation states, likely due to the alignment of latent manifolds during finetuning – a hypothesis supported by recent studies of representation learning. However, scGPT-HVG performance decreased from 14.28% to 6%, which can be attributed to the stricter filtering of genes used in scFME (compared to the zero-shot study), specifically silhouette score > 0.3.

Despite these improvements, both the fine-tuned Geneformer-30M and scGPT models remained below the performance of GEARS, a supervised deep learning model trained directly on perturbation data, which achieved 36.61% ISP accuracy as reported in [7]. It should be noted that scFME framework finetunes models to classify control vs. perturbed cells but not for predicting perturbed cell state. Whereas GEARS trains a deep learning model to predict perturbation directly. Therefore, these results suggest that while transfer learning from cell classification improves foundation model ISP performance, task-specific supervision retains an advantage for smaller models.

Notably, however, larger and domain-adapted foundation models outperformed GEARS. Geneformer-95M-Cancer, which incorporates extended pretraining on cancer-specific data, achieved 66.1% ISP accuracy—nearly doubling GEARS’ performance. This result underscores the value of scale and domain-specific pretraining in enhancing the biological fidelity of ISP predictions relative to task-specific supervised models.

Together these findings suggest that fine-tuning substantially improves the predictive power of foundation models for ISP tasks, but that model architecture, scale, and training data also remain critical determinants of performance.

### 2.7 Rank/K-Value analysis for target identification

While the statistically significant cosine-shift greater than the random baseline indicates that a model has successfully reproduced in-vivo perturbation of a ground truth gene in the correct direction, it does not inform us if random and unintentional in-silico perturbations of one or more genes is reproducing the ground truth gene’s in-vivo perturbation with even higher accuracy (i.e. “closer” in the latent space) than the ground truth gene. Such an occurrence represents poor specificity of a model’s ISP of a gene. To assess this aspect of in-silico perturbation (ISP) predictions, we used the Rank/K-value metric; a quantitative measure of how a ground truth gene is ranked relative to the random and unintentional gene perturbations, see Figure 5 and Table 3.

**Table 3.** Median, mean, and standard deviation of K-values for successful perturbations across models, along with the number and percentage of genes passing the random baseline test.

| Model | # ISP Genes<br>Passed Random Baseline | K |  |  |
| --- | --- | --- | --- | --- |
|  |  | Median | Mean | Standard Deviation |
| GF-30M | 56 (25%) | 1.5 | 12.5 | 17.38 |
| GF-95M | 126 (56.25%) | 6.5 | **12** | 14.53 |
| GF-95M-Cancer | 148 (66.1%) | 6 | 12.76 | 15.21 |
| scGPT-RS | 29 (12.95%) | 15 | 15.86 | 12.88 |
| scGPT-HVG | 6 (2.68%) | 20 | 14.6 | 12.62 |
| GenePT | 5 (2.23%) | 38 | 35.4 | 11.8 |

**Fig. 5.**
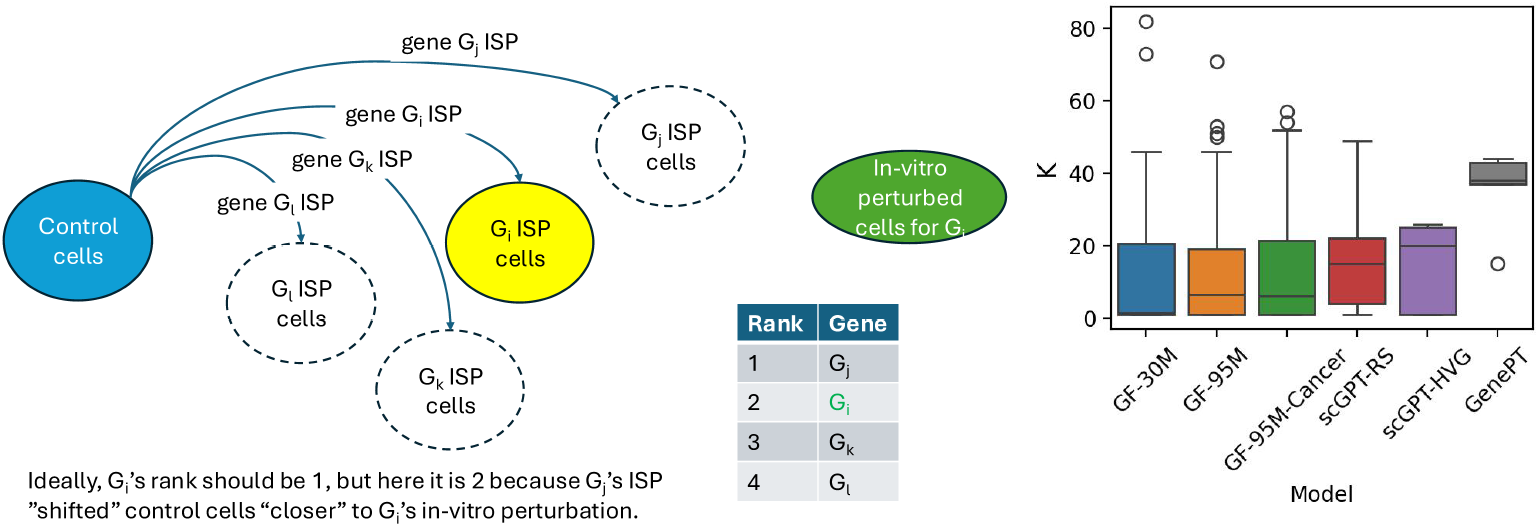
Distribution of K-values for successful perturbations across models. Lower K-values indicate higher specificity

For each successful ISP, we calculated the median cosine shift for all genes perturbed. Genes were ranked by median cosine shift (the highest cosine shift first), and the position of the ground truth gene in this ranking is denoted as K. A lower K value (higher cosine shift) indicates higher specificity, suggesting that the model shifts the control cell state toward the true target predominantly because of perturbation of a given ground truth gene, rather than by perturbation of unrelated genes by chance. The results show that Geneformer-30M scores the best among all the models, including the larger models

Across the evaluated models, Geneformer-30M achieved a median K of 1.5 for 56 (25%) successful perturbations, indicating strong specificity despite its smaller scale. Geneformer-95M and Geneformer-95M-Cancer, which incorporate larger training corpora and domain-specific data, yielded median K values of 6.5 and 6 respectively, with 126 (56.25%) and 148 (66.1%) successful perturbations. These results suggest that while model scale and biological relevance enhance overall ISP performance, they may also introduce unexpected perturbation effects, potentially reflecting increased sensitivity to transcriptomic context.

In contrast, scGPT-RS exhibited a median K of 15 across 29 (12.95%) successful perturbations, indicating lower specificity in ISP predictions.

These findings underscore the utility of the Rank/K-value metric in evaluating the specificity of ISP predictions. Models with lower K values are more likely to identify biologically relevant targets with high specificity, a critical consideration for therapeutic applications.

### 2.8 Biological function analysis of ISP performance

In this analysis, we used the scFME framework to highlight a model’s ISP performance on different gene categories based on biological functions and provide a comparison among the models in this aspect. First, gene set enrichment analysis on the ground truth genes was conducted using a gene-pathways resource, specifically STRINGdb [15]. This analysis revealed a strong biological connection for the 224 ground truth genes, showing significant enrichment (False Discovery Rate (FDR), < 0.05) of 23 terms from the GO (Gene Ontology) Molecular Function database.

Next, to evaluate whether any of the models exhibit biological functional bias in performance, we calculated sensitivity for these enriched terms containing more than 10 genes. A total of 191 genes out of 224 were assigned to at least one gene set. Figure 6 presents the sensitivity of each evaluated model per gene set; scGPT-HVG and GenePT were excluded due to the low number of successfully identified genes. Closely related gene sets were grouped into enriched classes based on gene set similarity (> 0.7) and overlap of enriched genes.

**Fig. 6.**
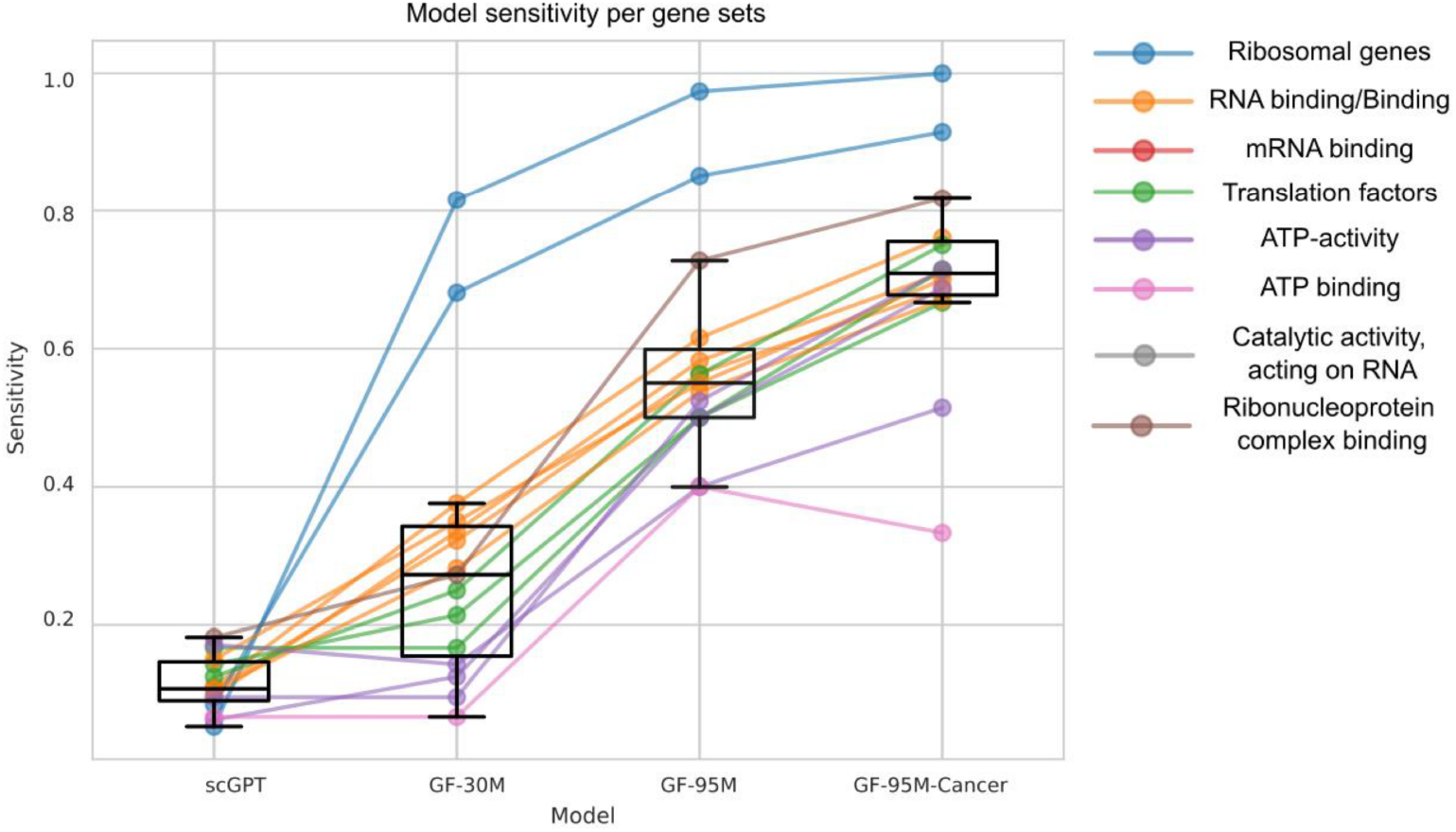
Sensitivity of ISP predictions for enriched gene sets grouped by functional classes. Geneformer models exhibit strong performance for ribosomal and translation-related genes, while scGPT shows relative strength in ATP-binding categories.

All models in the Geneformer family demonstrated exceptionally high sensitivity for ribosomal genes, which may be attributed to the high prevalence of ribosomal genes in the cell atlas used for pre-training the Geneformer models. Geneformer-30M and scGPT-RS models identified a similar total number of genes (56 vs. 29, see Table 1) but showed noticeable performance differences across enriched classes. scGPT-RS demonstrated better performance for ATP-activity/binding, comparable performance for translation factors, and significantly lower performance for binding and ribosomal genes compared to Geneformer-30M. We also observed a consistent improvement in sensitivity across all gene sets for Geneformer-95M compared to Geneformer-30M, and further improvement broadly across the gene sets for Geneformer-95M-Cancer compared to Geneformer-95M.

Interestingly, there were 32 genes that were not identified by any model. To assess whether there is any biological basis underlying these genes, we performed gene set enrichment using 224 genes as the background. This analysis resulted in no enriched gene sets, suggesting no biological reasoning behind these unsuccessful genes. This analysis shows that scFME framework can offer a systematic way to explore model ISP performance relative to gene functional categories.

## 3 Discussion

### Key findings

The scFME framework provides useful insights into ISP performance of single-cell foundation models, as demonstrated through application on seven distinct models. By leveraging three key metrics—the silhouette score, the number of successful gene perturbations relative to zero and random perturbation baselines, along with relevant statistical tests—the framework characterizes model performance in a robust manner. Notably, a high silhouette score, which reflects strong latent space separation between control and in-vitro perturbed cells, does not always correspond to superior ISP performance, as observed with the scGPT-RS model. Importantly, this approach allows for the assessment of model performance independently of models’ design criteria and leveraging manifold alignment though finetuning.

### Contribution and innovation

Unlike the zero-shot methods, this approach offers deeper insights by leveraging the manifold hypothesis, by fine tuning the models to train on the specific task. However, the framework does not require fine-tuning on perturbation data – finetuning for classification is adequate – and so the resulting model performance observations are likely to be relevant to scenarios where the datasets only contain the gene expression for disease and healthy cells.

### Generalizability

This approach is both simple and highly generalizable, allowing for the assessment of any model, regardless of whether it is fine-tunable. It can be applied to any perturbation dataset, provided that the number of cells and perturbed genes is sufficiently large. Fine-tuning need not be limited to classification tasks but can also encompass other objectives. Cosine similarity and shifts are versatile metrics that generalize well to models predicting gene expressions, such as encoder-decoder architectures, as well as to variational autoencoder (VAE) models that generate distributions rather than fixed embeddings.

### Biological implications

In-silico perturbation is increasingly recognized as a valuable downstream application of single-cell foundation models for target discovery and disease understanding. By incorporating ISP as a central component of the methodology, the framework directly addresses these significant biological implications. The inclusion of fine-tuning further enables the evaluation of a model’s capacity to discern the effects of perturbation sources, such as a disease scenario.

Applying the scFME framework to various foundation models revealed notable differences in ISP sensitivity across gene functional categories, likely reflecting distinctions in the latent cell and gene representations learned during pretraining and finetuning. Importantly, this framework also aids in the selection of models best suited for analyzing potential targets within specific gene classes, as it highlights differential model sensitivities across these categories.

Notably, the framework uncovered a potential connection between differential gene expression—measured by log fold change—and a model’s ability to successfully simulate cell-perturbation in silico. This finding could be important for a model’s ability to identify causal genes, when its expression change may be subtle relative to other genes, as seen in some realistic disease scenarios.

### Limitations

To ensure useful evaluation, it is necessary to have enough cells for fine-tuning and for conducting relevant statistical analyses, as well as an adequate number of gene perturbations to reliably estimate the random baseline. The framework attempts to align evaluation along the model’s manifold learning from fine tuning, therefore the model under evaluation should be capable of fine-tuning for the best model performance, in contrast to models like GenePT, where updating model embeddings through fine-tuning on the perturbation dataset is not straightforward. Additionally, a given model needs to be finetuned several times—corresponding to the number of ground truth genes—which can be time-consuming and resource-intensive.

### Future work

The current scFME framework suggests several future explorations of foundation models and benchmarking. It would be valuable to systematically compare models optimized via hyperparameter tuning, providing further insights into changes in their ISP performance. Additionally, focused investigation of each of the architectural differences between model families—i.e. Geneformer and scGPT—could clarify how design choices, such as input preprocessing, selection of pretraining data, cell representation, model architecture, contribute to ISP performance. On the evaluation front, analyzing on-target and off-target perturbation effects using Rank K-value metrics could yield deeper insights; it would be informative to determine which targets a gene’s ISP performance most closely approximates. Finally, more granular investigations at the gene level may uncover novel biological relationships, such as gene regulatory networks (GRN), thereby enriching our understanding of the interplay between model predictions and underlying biology.

## 4 Methods

### 4.1 Dataset and Preprocessing

The K562 Essential dataset from Weissman Labs was used for all benchmarking experiments. This single-cell Perturb-seq dataset comprises transcriptomic profiles of control and gene-perturbed cells and was obtained from Zenodo (record 13350497, version 1.4). Cells were filtered to retain those expressing at least 1,000 genes, with UMI counts between 1,000 and 100,000, and mitochondrial gene content below 15%. Doublets were removed. Genes were retained if expressed in at least 25 cells.

### 4.2 Ground Truth Selection

Ground truth genes were identified using a two-step filtering process:

#### 1. Differential Expression Analysis

Using the scanpy package (v1.10.1), we applied `scanpy.tl.rank_genes_groups` with a t-test to identify differentially expressed genes. Genes were retained if they showed a log2 fold change < − 1 and an adjusted p-value < 0.05. Cells were required to express at least 1,000 genes, have UMI counts between 1,000 and 100,000, and mitochondrial gene content below 15%. Doublets were removed, and genes had to be expressed in at least 25 cells.

#### 2. Transcriptomic Divergence

We applied the E-distance test using the scPerturb package (v0.1.0), with 10,000 runs and squared Euclidean distance as the metric. E-distance was chosen over silhouette score as we wanted to measure if in-vitro control and perturbed cells are significantly different and far way from each other, rather than well clustered. Equal subsampling to 200 cells was performed, and 2,000 highly variable genes were selected using the Seurat flavor. Genes with an adjusted p-value < 0.05 were retained.

E-distance can be defined as the difference in transcriptomic profiles between control and perturbed cells, quantifying how well a model can capture the perturbation effects. Let *X* = {*x*_1_, *x*_2_, …, *x*_*n*_} and *Y* = {*y*_1_, *y*_2_, …, *y*_*m*_} be two sets of vectors, *E* − *distance* can be computed as follows:

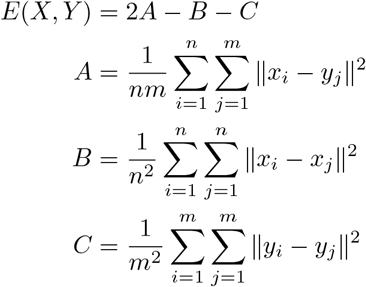

A total of 224 genes passed both filters in the K562 dataset and were used as ground truth for model fine-tuning and evaluation

### 4.3 Foundation models

We evaluated seven single-cell foundation models, each representing distinct architectural and pretraining strategies. All models were pretrained on large-scale single-cell RNA sequencing (scRNA-seq) datasets using self-supervised learning objectives and subsequently fine-tuned for the in-silico perturbation (ISP) task. GenePT is an exception on both counts. Default hyperparameter settings were used as recommended by the authors in GitHub or other relevant code repositories.

#### 4.3.1 Geneformer-30M

Geneformer-30M is a 6-layer Transformer model pretrained on 30 million healthy human cells using a masked language modeling (MLM) objective. Input sequences consist of the top 2,048 expressed genes per cell, ranked by normalized expression. During pretraining, 15% of gene tokens were masked and the model was trained to predict the masked genes based on surrounding context. This model serves as the baseline for the Geneformer family.

#### 4.3.2 Geneformer-95M

Geneformer-95M is a scaled-up variant of Geneformer-30M, featuring 12 Transformer layers and an extended input sequence length of 4,096 tokens. It was pretrained on 95 million cells, encompassing a broader range of biological contexts. The same MLM objective and ranked gene input strategy were used. The increased model capacity and input dimensionality are designed to enhance the model’s ability to capture complex transcriptomic patterns.

#### 4.3.3 Geneformer-95M-Cancer

This model variant builds upon Geneformer-95M by incorporating an additional 14 million cancer-specific cells into the pretraining corpus. The model architecture remains unchanged, but the inclusion of disease-relevant data is intended to improve performance on cancer-related perturbation tasks. This domain-adapted model enables evaluation of the impact of biologically aligned pretraining on ISP accuracy.

#### 4.3.4 scGPT-RS and scGPT-HVG

scGPT is a generative Transformer model pretrained on 33 million human cells using a generative masked modelling objective, akin to the GPT series. Unlike Geneformer, scGPT encodes gene expression as binned values rather than ranks. In the model we used, 51 bins were used. Input sequences vary in length depending on the task, with 3,000 gene tokens and 1 CLS used in this study. The model has demonstrated utility across multiple downstream tasks, including cell type annotation, batch correction, and perturbation prediction. We benchmarked two variants; the first, denoted as scGPT-RS (for random sample), allows the models to randomly select 3,000 non-zero expressed genes for each cell. The second, denoted as scGPT-HVG, identifies the top 3,000 most highly variable genes first prior to tokenizing the dataset. For each cell, non-zero expressed genes from these 3,000 highly variable genes are selected.

#### 4.3.5 GenePT

GenePT is a foundation model for single-cell biology that leverages large language model (LLM) embeddings—specifically OpenAI’s GPT-3.5—to generate biologically meaningful representations of genes and cells. Unlike traditional single-cell models that require extensive pretraining on raw gene expression atlases, GenePT uses curated textual descriptions of genes from the NCBI Gene database to construct gene-level embeddings. These embeddings are then aggregated to produce cell-level representations, enabling efficient and interpretable modeling of gene and cell properties across a wide range of downstream tasks.

### 4.4 Fine-tuning procedure

Each model was fine-tuned separately for each of the 224 ground truth genes to classify control versus perturbed cells. An 80/20 stratified train-test split was used. The majority class was not down sampled. Fine-tuning was not conducted for GenePT because it is not relevant to the model. Model-specific fine-tuning parameters were as follows:

- **Geneformer models:** learning rate = 1e–5, weight decay = 1e–3, batch size = 8, epochs = 20, linear learning rate scheduler, initial two layers were frozen for all variants.
- **scGPT models:** learning rate = 1e–4, batch size = 32, epochs = 20, 51 bins, 4 layers, 4 attention heads, dropout = 0.2, input length = 3,001.

### 4.5 Embedding Generation and Latent Space Analysis

For Geneformer and scGPT fine-tuned models, gene embeddings were extracted from the final encoder layers for each fine-tuned ground truth gene model. Cell embeddings were computed by averaging gene embeddings, excluding special tokens. For GenePT, weighted averaging was used to create cell embeddings, by multiplying the textual gene embeddings by the gene’s expression level. The mean of all cell embeddings was used as the reference point for the perturbed cell cluster of each ground truth gene in cosine similarity calculations. Silhouette scores were computed using `sklearn.metrics.silhouette_score` to assess separation between control and perturbed cells. Silhouette score is used to evaluate the quality of clustering, specifically cohesion and separation. Cohesion measures how close the point is to other points within the same cluster, and separation measures how far the point is from points in the nearest neighboring cluster. Values range from -1 to 1, where negative values typically indicate incorrect cluster assignments, values around 0 indicate overlapping clusters and values closer to 1 indicating well defined clusters. Silhouette score was used as a metric over E-distance as we want to measure how well clustered cell states are for ISP, rather than how far away the embeddings are in the latent space. Using trial-and-error exploration, we determined that 0.3 is a good threshold for models to distinguish well between control and perturbed cells. Average silhouette score can be calculated as follows for a given set of cells:

For a given sample *i*, define:

- *a*(*i*): the mean intra-cluster distance (average distance between *i* and all other points in the same cluster),
- *b*(*i*): the mean nearest-cluster distance (lowest average distance between *i* and all points in any other cluster).

Then, the silhouette coefficient for sample *i* is:

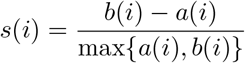

The average silhouette score over all samples is:

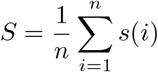

where *n* is the total number of samples.

### 4.6 In-silico perturbation (ISP)

ISP was simulated by modifying gene expression in control cells in the followings ways for each model:

- **Geneformer**: Target gene token moved to the end of the ranked list.
- **scGPT**: Binned expression set to zero.
- **GenePT**: Expression multiplied by the observed fold change from differential gene expression.

Perturbed gene embeddings were generated via forward passes through the fine-tuned models, and they were aggregated to form ISP cell embeddings. For GenePT, perturbed cell embeddings were generated directly by multiplying the textual gene embeddings with their perturbed gene expressions. Cosine similarities were computed between unperturbed and perturbed embeddings relative to the target state. The target state for each gene was derived by taking a mean of all cell embeddings for that given state. Cosine shift was defined as the difference between cosine similarity of the unpertured control cells and the target state, and the cosine similarity of perturbed cells and the target state. As cosine similarity ranges between [-1, 1], cosine shift ranges between [-2, 2].

- **Feature/Embedding space:** ℝ^*d*^ where d is the embedding dimension.
- **Target state:** *p* ∈ ℝ^*d*^. *P* is a set of target-state cell embeddings ∈ ℝ^*d*^, then:

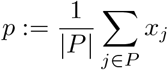
- **Control cell:** 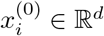
- **Perturbed cell (by gene** *g*): 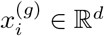.
- **Cosine similarity:**

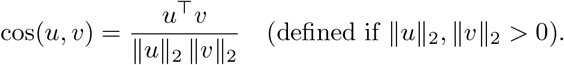

For a single control cell *i* and perturbation of gene *g*:

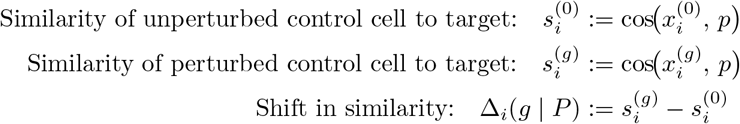

Interpretation: Δ_*i*_ > 0 means the perturbation moved cell *i* closer to the target state as measured by cosine similarity.

### 4.7 Statistical Analysis

Two statistical tests were applied:

#### 1. Zero-baseline test

One-sided Wilcoxon signed-rank test to assess whether cosine shift distribution as a result of in-silico perturbation of gene *g* over *n* control cells towards target state *P* is greater than 0.

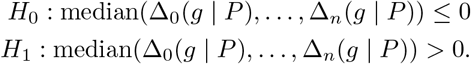

#### 2. Random baseline test

One-sided Wilcoxon rank-sum test to assess whether cosine shift distribution as a result of in-silico perturbation of gene *g* is greater than the cosine shift distribution of 100 randomly selected genes (*rand*) over *n* control cells towards target state *P*

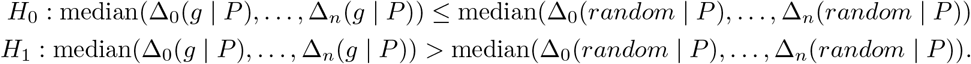

Multiple testing correction was performed using the Benjamini-Hochberg method (*α* = 0.05).

## 5 Conclusion

The key contribution of this work is the scFME framework, which offers a systematic and robust methodology for benchmarking fine-tuned foundation models for the insilico perturbation (ISP) downstream application, and demonstration of its application to various models to generate insights into their ISP performance. This framework ensures comprehensive assessment by requiring sufficient separation between control and perturbed cells and by quantifying ISP accuracy against random perturbations and zero-baseline references. Additionally, scFME enables detailed characterization of models’ performance relative to gene expression and gene functional categories.

The framework showed that larger models trained on larger datasets outperformed smaller models, supporting the scaling laws. The framework also highlighted the importance of fine-tuning in improving ISP accuracy, as seen in comparison to zero shot evaluation of the Geneformer and scGPT models. While further improvements are needed, the second generation Geneformer ISP performance is encouraging and shows considerable potential. Larger perturbation datasets are a critical need to study foundation models’ performance in further detail.

One primary direction for future research is the exploration of performance improvement with hyperparameter optimization using the framework, and the impact of larger perturbation datasets on the framework itself. Additionally, comparing the architectural differences between the Geneformer and scGPT model families using a generalized model, like TEDDY [16], that can be configured as one of these, with scFME could provide a controlled study into how cell representation choices impact ISP performance. Another important avenue for future work using scFME is further exploration of models’ performance relative to various gene categories and relative expression levels of causal genes.

In conclusion, the scFME framework lays a strong foundation for future research and development in the evaluation of single-cell foundation models for in-silico perturbation. By building on the insights and advancements presented here, researchers and practitioners can continue to push the boundaries of what is possible and drive progress in this highly dynamic area.

## References

[1] Theodoris, C.V., Xiao, L., Chopra, A., Chaffin, M.D., Al Sayed, Z.R., Hill, M.C., Mantineo, H., Brydon, E.M., Zeng, Z., Liu, X.S., Ellinor, P.T.: Transfer learning enables predictions in network biology. Nature 618(7965), 616–624 (2023) 10.1038/s41586-023-06139-9

[2] Cui, H., Wang, C., Maan, H., Pang, K., Luo, F., Duan, N., Wang, B.: scgpt: toward building a foundation model for single-cell multi-omics using generative ai. Nature Methods 21(8), 1470–1480 (2024) 10.1038/s41592-024-02201-0

[3] Luecken, M.D., Gigante, S., Burkhardt, D.B., Cannoodt, R., Strobl, D.C., Markov, N.S., Zappia, L., Palla, G., Lewis, W., Dimitrov, D., Vinyard, M.E., Magruder, D.S., Mueller, M.F., Andersson, A., Dann, E., Qin, Q., Otto, D.J., Klein, M., Botvinnik, O.B., Deconinck, L., Waldrant, K., Yasa, S.N., Szalata, A., Benz, A., Li, Z., Rieck, B., Ahlmann-Eltze, C., Veiga Beltrame, E., Bravo González-Blas, C., Chen, A.T., DeMeo, B., Ergen, C., Floc’hlay, S., Gayoso, A., Hicks, S., Ji, Y., Kleshchevnikov, V., La Manno, G., Lombardo, M.G., Lopez, R., Righelli, D., Sarkar, H., Svensson, V., Tong, A., Xing, G., Xu, C., Bloom, J.M., Pisco, A.O., Saez-Rodriguez, J., Wulsin, D., Pinello, L., Saeys, Y., Theis, F.J., Krishnaswamy, S., Members, O.P.J.: Defining and benchmarking open problems in single-cell analysis. Nature Biotechnology 43(7), 1035–1040 (2025) 10.1038/s41587-025-02694-w

[4] Brooks, T.G., Lahens, N.F., Mrčela, A., Grant, G.R.: Challenges and best practices in omics benchmarking. Nature Reviews Genetics 25(5), 326–339 (2024) 10.1038/s41576-023-00679-6

[5] Buchka, S., Hapfelmeier, A., Gardner, P.P., Wilson, R., Boulesteix, A.-L.: On the optimistic performance evaluation of newly introduced bioinformatic methods. Genome Biology 22(1), 152 (2021) 10.1186/s13059-021-02365-4

[6] Ahlmann-Eltze, C., Huber, W., Anders, S.: Deep learning-based predictions of gene perturbation effects do not yet outperform simple linear baselines. bioRxiv (2025) 10.1101/2024.09.16.613342 https://www.biorxiv.org/content/early/2025/02/07/2024.09.16.613342.full.pdf

[7] Liu, X., Boylan, J., Bouiller, T., Solovyeva, E., Ataman, B., Hoersch, S., Jenkins, J., Devarakonda, M.: Evaluating foundation models for insilico perturbation. bioRxiv (2025) 10.1101/2025.05.11.653338 https://www.biorxiv.org/content/early/2025/05/15/2025.05.11.653338.full.pdf

[8] Loaiza-Ganem, G., Ross, B.L., Hosseinzadeh, R., Caterini, A.L., Cresswell, J.C.: Deep generative models through the lens of the manifold hypothesis: A survey and new connections (2024) 2404.02954 [cs.LG]

[9] Gilpin, W.: The cell as a token: high-dimensional geometry in language models and cell embeddings. arXiv 2503.20278 (2025)

[10] Fang, Y., Ohn, I., Gupta, V., Lin, L.: Intrinsic and extrinsic deep learning on manifolds (2023). https://arxiv.org/abs/2302.08606

[11] Kojima, T., Gu, S.S., Reid, M., Matsuo, Y., Iwasawa, Y.: Large Language Models are Zero-Shot Reasoners (2022). http://arxiv.org/abs/2205.11916

[12] Dixit, A., Parnas, O., Li, B., Chen, J., Fulco, C.P., Jerby-Arnon, L., Marjanovic, N.D., Dionne, D., Burks, T., Raychowdhury, R., Adamson, B., Norman, T.M., Lander, E.S., Weissman, J.S., Friedman, N., Regev, A.: Perturb-seq: Dissecting molecular circuits with scalable single-cell rna profiling of pooled genetic screens. Cell 167(7), 1853–186617 (2016) 10.1016/j.cell.2016.11.038

[13] Peidli, S., Green, T.D., Shen, C., Gross, T., Min, J., Garda, S., Yuan, B., Schumacher, L.J., Taylor-King, J.P., Marks, D.S., Luna, A., Blüthgen, N., Sander, C.: scperturb: harmonized single-cell perturbation data. Nature Methods 21(3), 531–540 (2024) 10.1038/s41592-023-02144-y

[14] Xiao, Y., Hsiao, T.H., Suresh, U., Chen, H.I., Wu, X., Wolf, S.E., Chen, Y.: A novel significance score for gene selection and ranking. Bioinformatics 30(6), 801–807 (2014) 10.1093/bioinformatics/btr671 . Epub 2012 Feb 9

[15] Szklarczyk, D., Kirsch, R., Koutrouli, M., Nastou, K., Mehryary, F., Hachilif, R., Gable, A.L., Fang, T., Doncheva, N.T., Pyysalo, S., Bork, P., Jensen, L.J., von Mering, C.: The string database in 2023: protein-protein association networks and functional enrichment analyses for any sequenced genome of interest. Nucleic Acids Research (Database issue) 51 (2023)

[16] Chevalier, A., Ghosh, S., Awasthi, U., Watkins, J., Bieniewska, J., Mitrea, N., Kotova, O., Shkura, K., Noble, A., Steinbaugh, M., Delile, J., Meier, C., Zhukov, L., Khalil, I., Mukherjee, S., Mueller, J.: TEDDY: A Family Of Foundation Models For Understanding Single Cell Biology (2025)

